# Insights into the molecular features of the von Hippel-Lindau like protein

**DOI:** 10.1101/407353

**Authors:** Federica Quaglia, Giovanni Minervini, Francesco Tabaro, Silvio C.E. Tosatto

## Abstract

We present an *in silico* characterization of the von Hippel-Lindau like protein (VLP), the only known human paralogue of the von Hippel-Lindau tumor suppressor protein (pVHL). Phylogenetic investigation showed VLP to be mostly conserved in upper mammals and specifically expressed in brain and testis. Structural analysis and molecular dynamics simulation show VLP to be very similar to pVHL three-dimensional organization and binding dynamics. In particular, conservation of elements at the protein interfaces suggests VLP to be a functional pVHL homolog potentially possessing multiple functions beyond HIF-1α-dependent binding activity. Our findings show that VLP may share at least seven interactors with pVHL, suggesting novel functional roles for this under-studied human protein. These may occur at precise hypoxia levels where functional overlap with pVHL may permit a finer modulation of pVHL functions.

## Introduction

Cellular specialization arises from finely tuned interactions between different molecular components, such as proteins, DNA, RNA, transcription factors, degradative machinery and external factors [1]. This complex organization allows coordination of several metabolic networks yielding cellular metabolism. Oxygen sensing is an example of a biological network aimed at promoting metabolic adaptation to environmental variations, as is a well-characterized molecular pathway triggering complex responses to hypoxia [2,3]. Oxygen variation is perceived by the cell through interactions of highly specialized proteins: the von Hippel-Lindau tumor suppressor protein (pVHL), Hypoxia Inducible Factor 1α (HIF-1α) and prolyl-hydroxylase protein family (PHDs), which form the so-called core hypoxia response network [4]. At physiological oxygen concentrations, PHDs catalyse the hydroxylation of two specific HIF-1α prolines (P402 and P564) [5,6]. Hydroxylated HIF-1α is then targeted for proteasomal degradation by pVHL, an E3 ubiquitin ligase complex substrate recognition element [7]. Hypoxia inhibits PHD activity, impairing HIF-1α and pVHL recognition, promoting HIF-1α stabilization [8]. Once stabilized, HIF-1α is translocated to the nucleus, where it activates Hypoxia response elements (HRE) and promotes hypoxia-regulated gene expression [9]. Deregulation of this network is known to predispose to cancer onset, as in von Hippel-Lindau syndrome [10]. Biological networks, beyond their complexity, are also characterized by robustness [11], a fundamental property which allows the cell to react to internal and external perturbations (e.g. hypoxia, nutrient variation, DNA mutations, etc.) [11]. A bow-tie architecture is frequently used to describe biological networks [11,12]. This representation implies few highly connected hub proteins forming a conserved core, while a number of proteins and various stimuli form the network input/output. Elements in the conserved core are associated with the main tumor suppressors and oncogenes, such as p53 [13,14], pVHL [15] and c-Myc [16] just to name a few. Another property directly derived from a bow-tie network organization is the appearance of specialized protein hub isoforms (or alternative genes) presenting slightly different functions. The p53-like protein family is an example of this functional overlap within members of the same family [17]. p53, p63 and p73 are known to form a transcription factor family [18], acting as main regulators of cell cycle progression. Although partially redundant, each member has its own unique function, varying between coordination of cell differentiation, stress response, functional apoptosis and aging [17]. From a biological network perspective, functional overlap can be seen as a natural mechanism to improve network robustness, i.e. together replicating crucial nodes while allowing evolution of new functions. In 2004, the von Hippel-Lindau like protein (VLP), a human paralog of the VHL gene, was reported in the literature [19]. VLP contains 139 amino acids, corresponding to the pVHL β-domain, while the pVHL E3-component recognition domain, required for pVHL degradative function, is missing. The first characterization attempts revealed that VLP binds HIF-1α and suggested it may act as a dominant-negative VHL, protecting HIF-1α from degradation [19]. Three different pVHL isoforms [20] are already known in the literature. Although pVHL was described to be directly involved in several different pathways interacting with a number of different interactors [15], the exact role of each different isoform and paralogous VLP is far from being clearly understood. Here, we present an *in silico* characterization of human VLP. Phylogenetic analysis was paired with *in silico* structural investigations. Our analysis showed that VLP is expressed in specific human tissues and shares interactors with pVHL. Collectively, our findings suggest novel functional roles for this human pVHL paralog.

## Materials and Methods

### Expression data

Expression profile data was retrieved by combining different database searches. Bgee [21] was used to extract and compare gene expression patterns between homologous genes from different species and CleanEx [22] to extract specific entries according to their biological annotation. Ensembl [23], GeneCards [24], NextProt [25], GTEx Portal [26] and Gene Expression Atlas [27] were used to retrieve human gene expression data at the tissue level as well as to extract data for gene regulation and genetic variation.

### Sequence feature and interactor analysis

Orthologous VLP sequences from different species were retrieved in OMA Browser [28], OrthoDB [29], EggNOG [30], PhylomeDB [31] and enriched by manual Blast [32] search against the NR database. The extracted dataset was then manually curated to remove pVHL-specific homologous sequences and aligned with Jalview [33] to derive a conservation score for each residue. The final conservation score and chemical properties were considered as a cutoff for protein-protein interaction surface prediction. A VLP interaction network was derived merging data from the BIOGRID [34], STRING [35] and IntAct [36]. Putative VPL molecular functions were predicted with INGA [37], while dSysMap [38], COSMIC [39] and PolyPhen-2 [40] were used to address VLP role in human disease onset.

### Phylogenetic analysis

A bayesian analysis was performed with MrBayes 3.2.6 [41] using the Jones-Taylor-Thornton model of protein evolution plus invariant sites, JTT + I, identified with MEGA 7 [42], with 4 chains (3 heated chains and 1 cold chain) and temperature 0.1 and 1,000,000 generations. The tree layout and bootstrap values at the nodes of the tree were visually inspected with FigTree (http://tree.bio.ed.ac.uk/).

### Homology modeling

The crystal structure of pVHL bound to the CODD fragment of HIF-1α (PDB code: 1LM8) was selected as a template to model the VLP structure. The human VLP sequence (accession code: Q6RSH7) was retrieved from UniProt [43] and aligned with T-Coffee [44] against pVHL using standard parameters. This was used for model building in Modeller [45]. The VLP-HIF-1α dimer was modeled through structural superimposition in Chimera [46]. GROMACS 4.6.5 [47] was used to relax the model structures for 1.000 steps of steepest descent minimization. Model quality was assessed with Qmean [48] and TAP [49]. A network of interacting residues was generated using RING 2.0 [50] and used to inspect the main structural feature of VLP.

### Molecular dynamics simulation

Molecular dynamics (MD) simulations were carried out with Gromacs 4.6.5, using the CHARMM-22 force field [51] on a standard x86 Linux workstation. Hydrogen atoms were added to the system using the Gromacs pdb2gmx routine. The simulation protocol consists in a minimization step, 100 ps of NVT (constant number of molecules, volume and temperature) simulation, 100 ps of NPT (constant number of molecules, pressure and temperature) simulation and 50 ns of classical MD. Temperature for production simulations was kept at 300 K. A solvent box was generated imposing a cut-off distance of 10 Å from the farthest point of the protein boundaries in each dimension. Parameters for simulating the hydroxyproline residue were derived from [52].

## Results

The human VLP is a 139 amino acid protein with 67.6% identity and 78% similarity to the pVHL region 1–157. This corresponds to the N-terminus and β-domain of pVHL [53]. VLP proposed to co-regulate HIF-1α stability [19], however, to date its molecular function is far from understood. The results from different *in silico* analyses aimed at characterizing VLP are discussed in the following.

### VLP is found in cerebellum, testis and placenta

The expression patterns associated to VLP were investigated by a virtual expression analysis searching different databases. Multiple database searches in Bgee, CleanEX and Gene Atlas confirmed the previous observations reporting the human VLP in cerebellum and placenta [19]. The Gene Cards and GTEx Portal report VLP to be also expressed in testis (Figure 1). As these databases use data from different sources, this data cannot be considered a spurious observation. In particular, GTEx Portal collects novel analyses of human tissues from donors; it could be supposed that VLP expression could be caught in the near future within other tissues beyond brain cerebellum, placenta and testis. Of note, all of these tissues present a well known susceptibility to hypoxia [54–56]. Due to its molecular similarity with pVHL, a tissue-specific role for VLP could be hypothesized.

**Figure 1.**
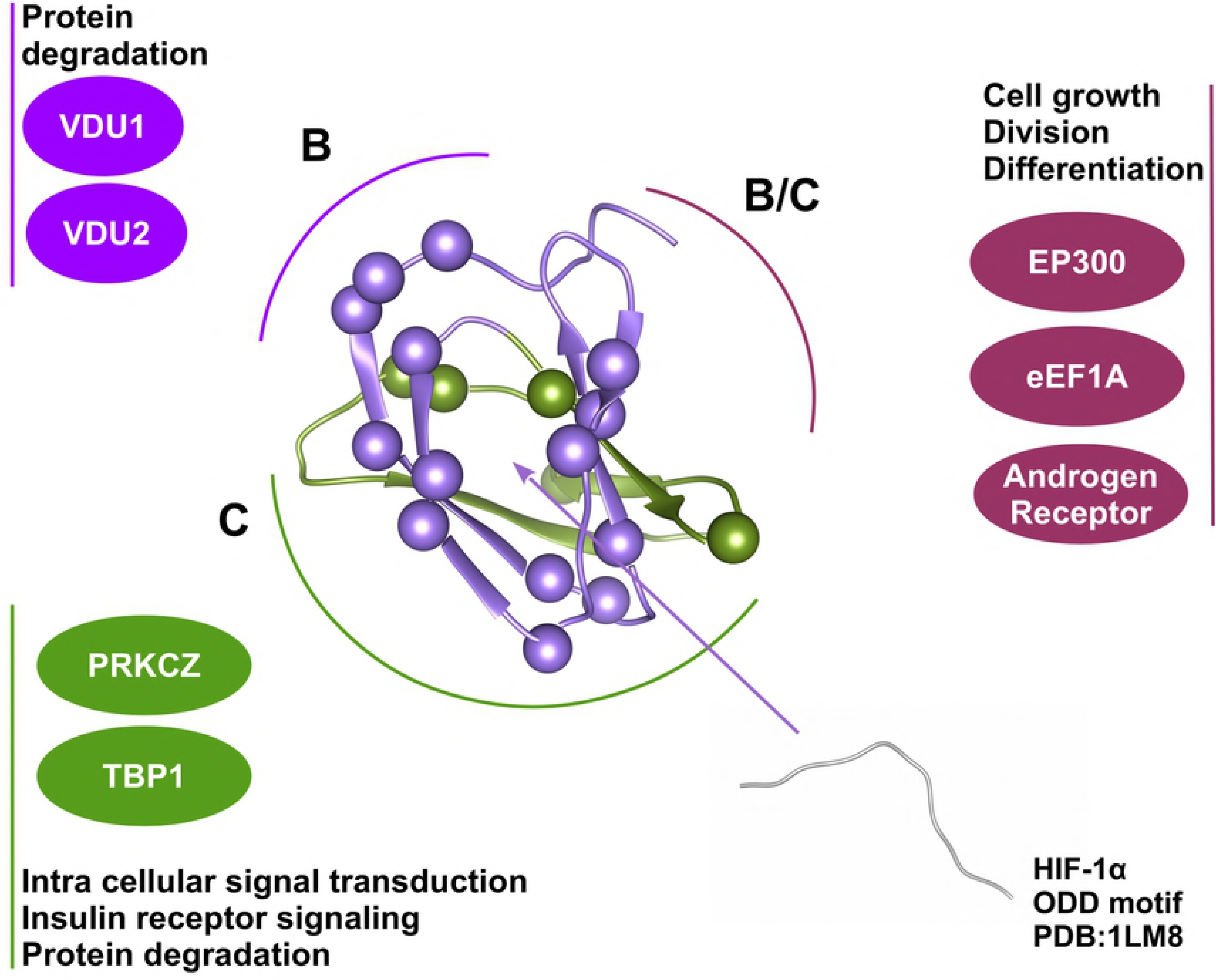
VLP sequence conservation. Multiple sequence alignment of VLP showing relevant conservation for the globular domain corresponding to the β-domain of pVHL. Blue box for the α-domain of pVHL missing in VLP. Color bars represent putative protein-protein interaction interfaces predicted for VLP using pVHL description as reported in [53]. VLP cancer mutations found in COSMIC are shown with red dots.

### VLP is mainly conserved among primates

Next, we asked whether VLP is conserved among other species. To answer this question, a comparative sequences analysis was done. Retrieving orthologous sequences is particularly challenging due to high sequence identity with the pVHL protein. Although an initial search using an automatic procedure generated a list of 47 putative orthologs, manual inspection showed that only a limited number of sequences can be correctly classified as VLP. In particular, VLP is mainly present in primates (*Homo sapiens*, *Pan paniscus*, *Nomascus leucogenys*, *Aotus nancymaae*, *Rhinopithecus roxellana*, *Mandrillus leucophaeus*), with *Vicugna pacos* and *Monodelphis domestica (the grey short-tailed opossum*, *a marsupial*) representing the only two other mammals whose sequences are not ambiguous and clearly orthologous to human VLP (Figure 2). A multiple-sequence alignment comparing a total of 7 VHL orthologs from distinct upper mammals shows the VLP globular region, corresponding to the β-domain of pVHL, to be highly conserved (> 85% similarity). A longer primate-specific region formed by three repetitions of the xEEx motif was found at the N-terminus (Figure 2). This sequence organization is comparable to pVHL30, the longest human pVHL isoform [57] which presents a 54 residue tail formed by five GxEEx repetitions. Although debated, this tail was proposed to mediate isoform-specific functions presumed to have emerged during primate evolution, i.e. a further protein-protein interaction interface [57].

**Figure 2.**
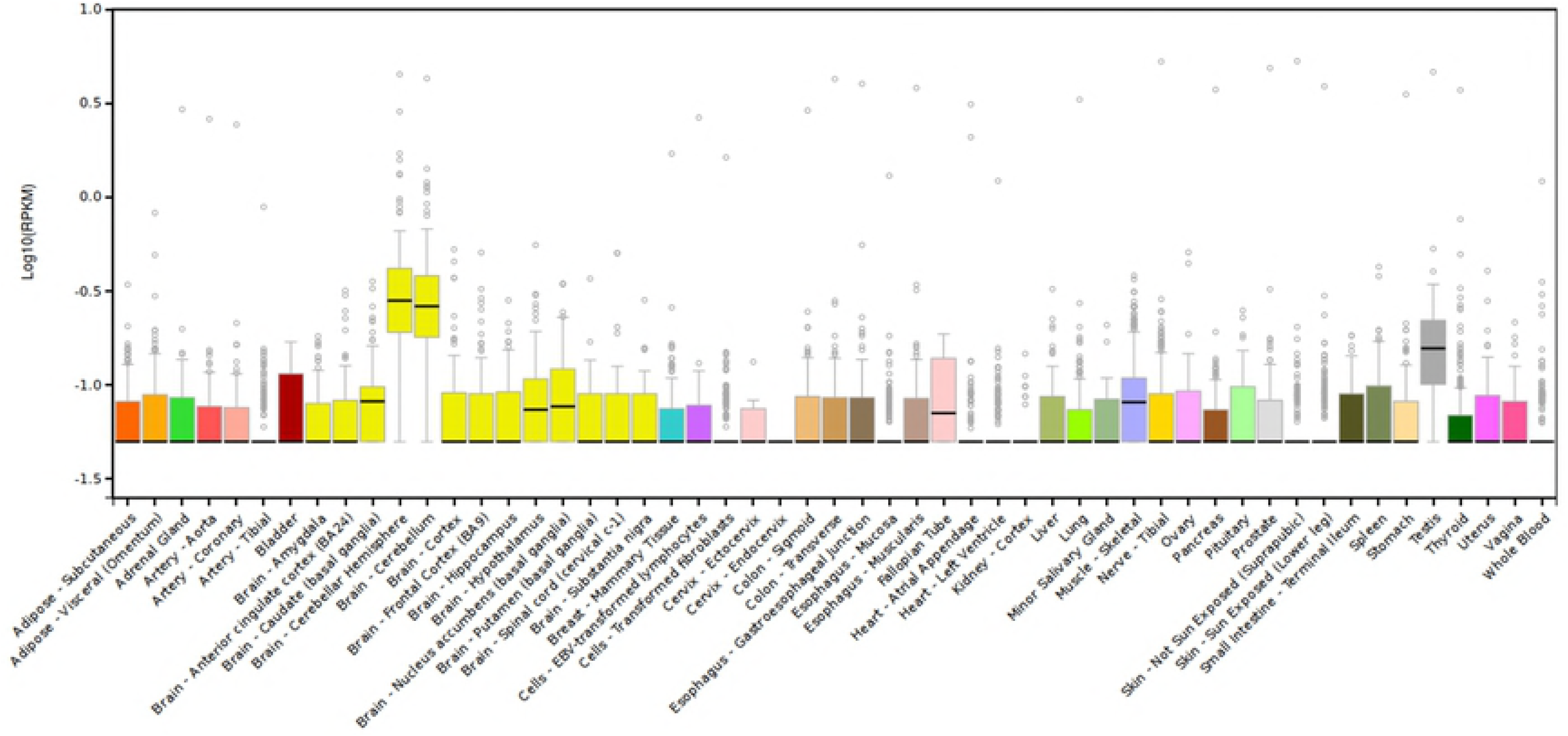
VLP expression in human. Box plot of human VLP gene expression data from GTEx Portal. VLP is actively expressed in testis and brain (cerebellum and basal ganglia).

### VLP phylogenetic analysis

Despite great efforts in retrieving orthologous sequences of human VLP from various sources, multiple sequence alignment suggested that not all “VHL-like proteins” are truly orthologous to human VLP. Indeed, most of them share a sequence and domain composition resembling VHL rather than human VLP. Due to this ambiguity, we discarded these putative sequences from the following analysis. We next performed a Bayesian inference phylogenetic analysis to better investigate the evolutionary relationships between VLP, its paralog VHL, and their different functions. Our phylogeny suggests a complex evolutionary relationship behind VLP and its paralog VHL. A group of primate VHL sequences can be identified, including *Homo sapiens*, while the VLP and VHL proteins from *Vicugna pacos* seem to form a separated group, sharing higher sequence similarity with the VHL group (Figure 3). As the Alpaca (*Vicugna pacos*) is a camelid evolved to live at higher altitudes of the South American mountains, the observed differences may reflect an adaptation to constitutive low oxygen concentrations. The analysis also highlights a well defined group including all VLP sequences from Primates, suborder Haplorhini, with a clear distinction for each family: Hominidae, including *Homo sapiens*, *Gorilla gorilla*, *Pan paniscus* and *Pongo abelii*, and Hylobatidae represented by *Nomascus leucogenys*. On the other hand, we found no trivial localization for *Monodelphis domestica* as its putative VLP shares features with both VHL and VLP groups. These findings, paired with the high sequence similarity shared by the VHL *β*-domain and VLP sequences, suggest that one (or more) remote gene duplications during mammalian evolution could easily explain the differences observed between VLP sequences from distant groups. Further genetics characterization will help to clarify this point.

**Figure 3.**
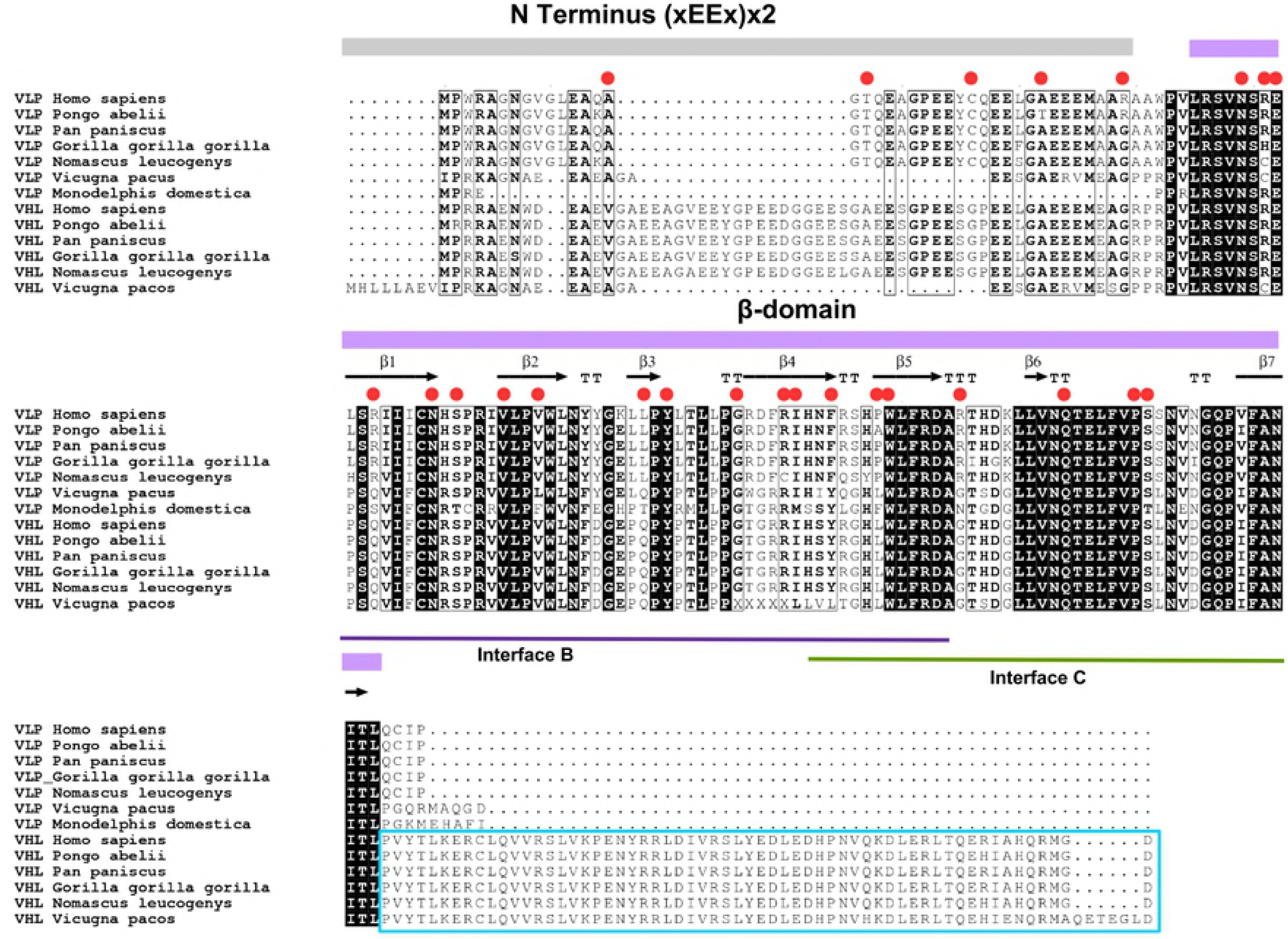
Phylogenetic analysis of VLP and pVHL. The VLP group is highlighted in orange, with sky-blue box used for pVHL. Bootstrap values are shown at nodes.

### VLP presents a structured *β*-domain adorned by a long disordered tail

A VLP homology model, generated starting from pVHL structure (PDB code 1LM8 [58]), shows a clear pVHL *β*-domain fold, completely lacking the elongin binding α-domain with a long disordered N-terminus (Figure 4). Superimposition of the VLP model and pVHL crystal structure highlights full conservation (100% identity) for the *β*2- and *β*6-strands and a short loop in pVHL. The model quality calculated with QMEAN is 0.649, while TAP estimates a 0.7569 confidence value, suggesting that our VLP model is good. Lower reliability was estimated for the N-terminal tail. MobiDB [59] confirmed the disordered behavior of the region. We then decided to perform molecular dynamics simulations to verify VLP fold stability. A 50 ns MD simulation showed that the model is stable (average RMSD 1.8 Å and 2.5 Å for the globular domain and entire protein, respectively). Of note, transient short α-helix segments were noted during simulations in the disordered tail, suggesting that this region is dynamically active and prone to explore different conformations.

**Figure 4.**
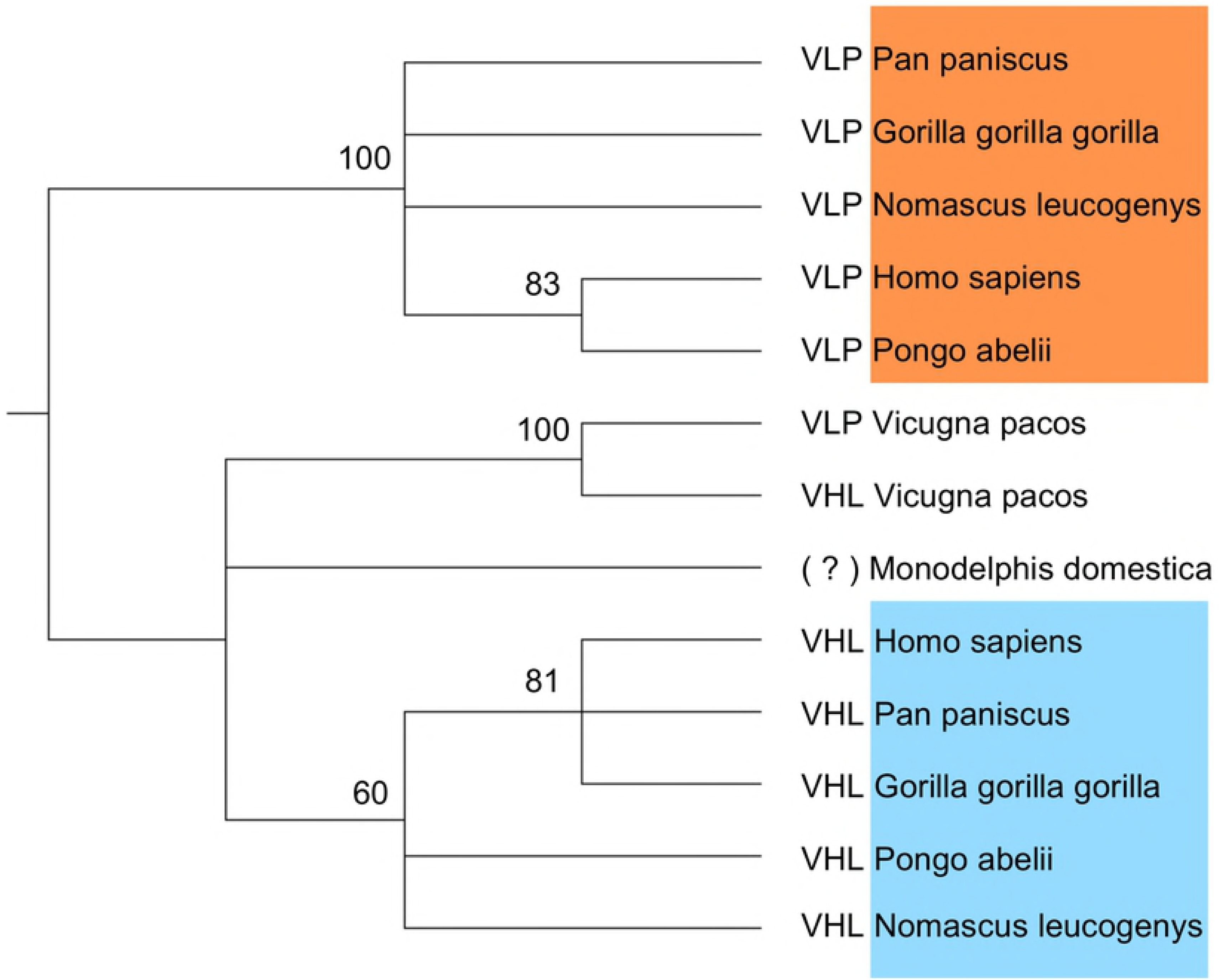
Overview of VLP structural features. Structural comparison of domain organization and electrostatic surface between VLP and pVHL. VLP homology model in ribbon representation with semitransparent surface colored by conservation (purple, high; turquoise, low). VLP corresponds to *β*-domain of pVHL (blue box).

### VLP interacts with HIF-1α through interface B

Although interaction between VLP and HIF-1α is not completely new, driving molecular details remain unknown. Due to its similarity with pVHL the interaction was proposed to rely on similar residues [19]. Our structural model shows amino acids substitutions at the putative HIF-1α interaction interface are present which may interfere with VLP/HIF-1α interaction. The corresponding VLP complex was modeled by superimposition with the pVHL/HIF-1α complex crystal structure. Two independent MD simulations were performed to test complex stability with and without HIF-1α hydroxylation (Figure 5). These showed VLP interacting with HIF-1α through the *β*3- and *β*4 strands. The binding is maintained by electrostatic interactions between VLP residues and the HIF-1α region comprising residues 561-565. The remaining residues appear to follow a different fate depending on presence of hydroxylated proline 564. Simulations showed that hydroxylation of proline 564 may play a regulative role in the interaction with VLP. The hydroxylated complex is more stable during the simulation, in particular a hydrogen bound between hydroxyproline 564 and histidine 97 of VLP lower the average distance between the two proteins, yielding further hydrophobic interactions with interface B residues (Figure 5).

**Figure 5.**
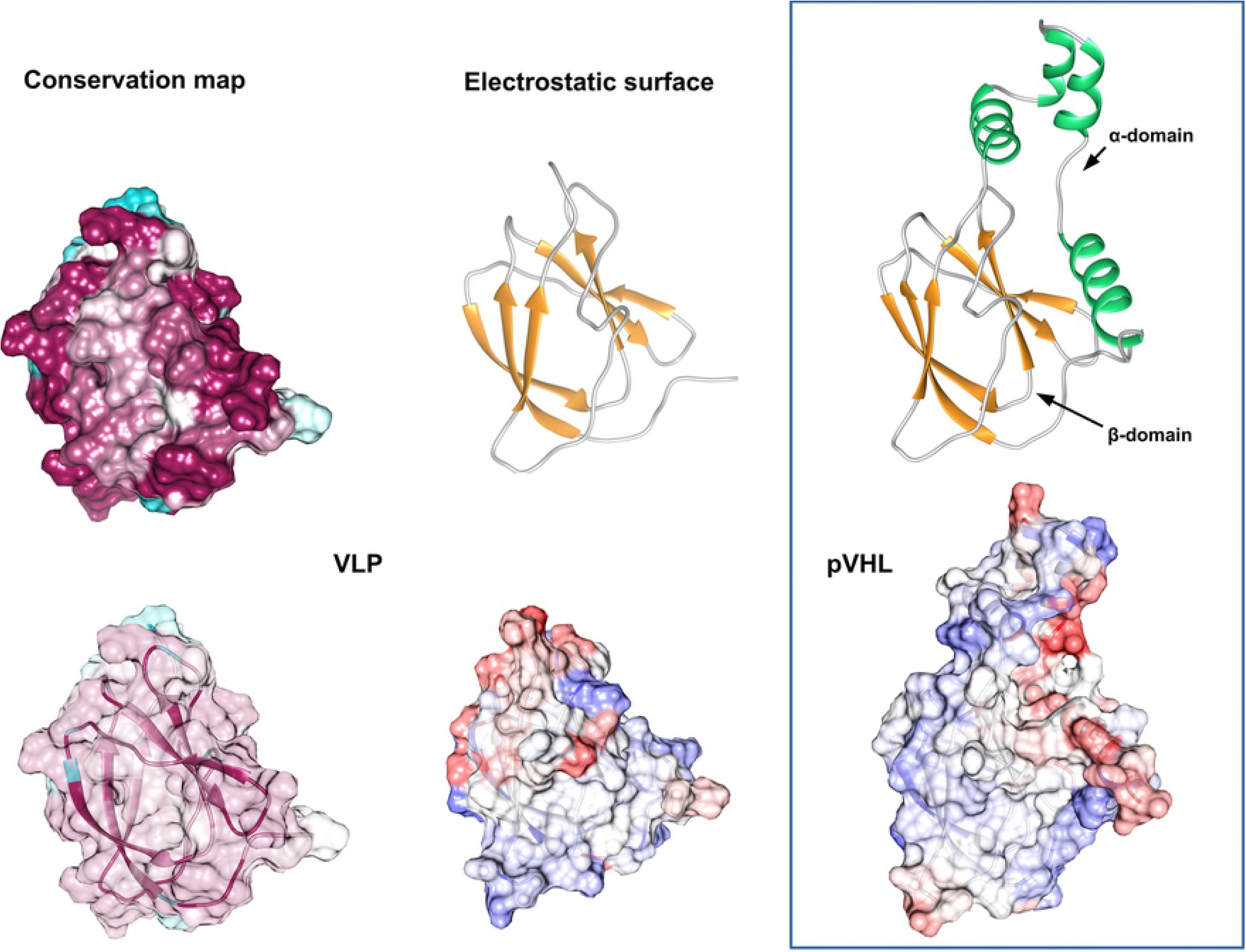
Molecular dynamics simulation of VLP/HIF-1α interaction and VLP mutation analysis. The starting VLP/HIF-1α complex is shown on the left (blue box). The results of 40 ns of simulation for wild type HIF-1α (top right, red box) revealed VLP forms stable interaction with the LxxLAP motif [68], while presence of hydroxylation (bottom right, green box) reinforces the interaction yielding enlarged interaction surface. The HIF-1α ODD motif is shown in light blue and red is used for VLP interacting residues. Light grey is for region of VLP not involved in the binding.

### VLP presents conserved structural motifs suggesting overlapping pVHL functions

In a previous research, we proposed three different pVHL protein-protein interaction interfaces (A, B and C, respectively) [53]. For each interface, conserved residues are known to engage in specific interactions with a subset of pVHL binding partners [60]. We asked whether a similar surface organization is also conserved in VLP. Considering the high sequence identity between the two proteins, we mapped putative binding regions on the VLP interface. Residues forming interfaces B and C are mostly conserved in VLP. Considering the similarity between the two proteins, at least seven pVHL interactors can be inferred for VLP (Figure 6). Putative pathways in which VLP may play a role were assigned based on these findings. VLP may be involved in regulating HIF-1α degradation, cell migration and cell cycle regulation interacting with VDU1/USP30 and VDU2/USP20 through interface B. Both are well known deubiquitinating enzymes involved in various cellular processes [61–63]. For interface C, interactions with EP300, eEF1A, Androgen receptor and TBP1 were predicted. This suggests VLP to be involved in various pathways, such as cell cycle regulation, nuclear export, growth and development of male reproductive organs and ATP-dependent degradation of ubiquitinated proteins.

**Figure 6.**
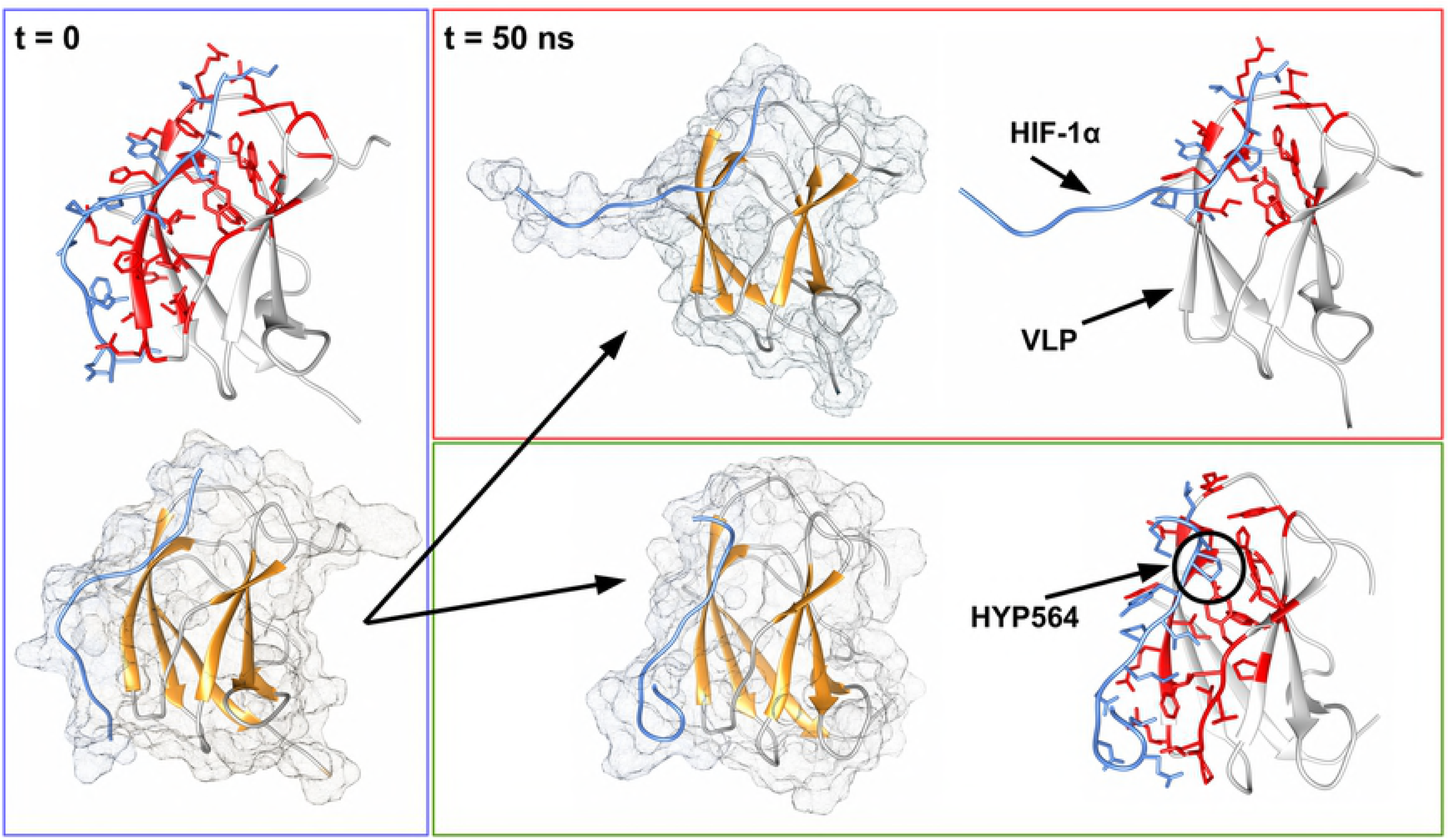
VLP in the cellular context. Conservation analysis suggests 7 putative interactors shared with pVHL beyond HIF-1α. Interactions rely on the putative interaction interfaces B and C highlighted in the structure as purple and green, respectively. Interactors associating multiple interface are presented as wine-colored. Putative functional pathways are presented above and associate to each interface. Spheres represent VLP mutation sites and are colored according to the affected VLP binding interfaces.

### VLP mutations

COSMIC contains 37 mutated VLP samples, including 26 missense, 9 synonymous, 1 nonsense substitution and 1 non-stop gain substitutions. No other complex mutations were found. Both confirmed somatic mutations and variants of unknown origin are described. A schematic view of the distribution of these mutations on the VLP sequence and their pathogenicity predictions are presented in Table 1. VLP mutations appear to uniformly affect the entire protein. Interestingly, tissues and organs affected are largely different from those affected in the VHL syndrome (Figure 7). Considering only missense mutations, 33.6% were found in adenocarcinoma of the large intestine, 15.4% in gastric adenocarcinoma and 11.5% in endometrioid carcinoma. Others organs affected include biliary tract, breast, small intestine, kidney, liver, lung, ovary, esophagus and skin.

**Table 1.** VLP mutations and pathogenicity prediction. For each VLP mutation found in COSMIC, we report the amino acid position and type for both wild type and mutated residues. Conservation in sequence and PolyPhen predictions (benign, possibly damaging or probably damaging) are also shown. “n/a” is used for non-missense mutations.

| Position | Wild type | Mutant | Conserved | PolyPhen 2 |
| --- | --- | --- | --- | --- |
| 15 | Ala | Val | yes | possibly damaging |
| 17 | Thr | Ala | yes | benign |
| 26 | Cys | Tyr | yes | possibly damaging |
| 32 | Ala | Val | yes | benign |
| 39 | Arg | Gly | no | benign |
| 49 | Asn | Lys | yes | probably damaging |
| 51 | Arg | Cys | no | benign |
| 52 | Glu | Gly | yes | probably damaging |
| 55 | Arg | Trp | no | probably damaging |
| 60 | Asn | Lys | yes | probably damaging |
| 62 | Ser | Asn | yes | probably damaging |
| 66 | Val | Met | yes | probably damaging |
| 69 | Val | Leu | yes | benign |
| 70 | Trp | - | yes | n/a |
| 78 | Leu | Met | no | probably damaging |
| 80 | Tyr | Asp | yes | probably damaging |
| 86 | Gly | Ser | yes | probably damaging |
| 90 | Arg | His | no | possibly damaging |
| 91 | Ile | Thr | yes | benign |
| 94 | Phe | Leu | yes | possibly damaging |
| 98 | Pro | His | yes | probably damaging |
| 99 | Trp | Arg | yes | probably damaging |
| 105 | Arg | Ser | yes | benign |
| 114 | Gln | Lys | yes | possibly damaging |
| 120 | Pro | Leu | yes | probably damaging |
| 121 | Ser | Tyr | yes | probably damaging |
| 140 | - | Arg | - | n/a |

**Figure 7.**
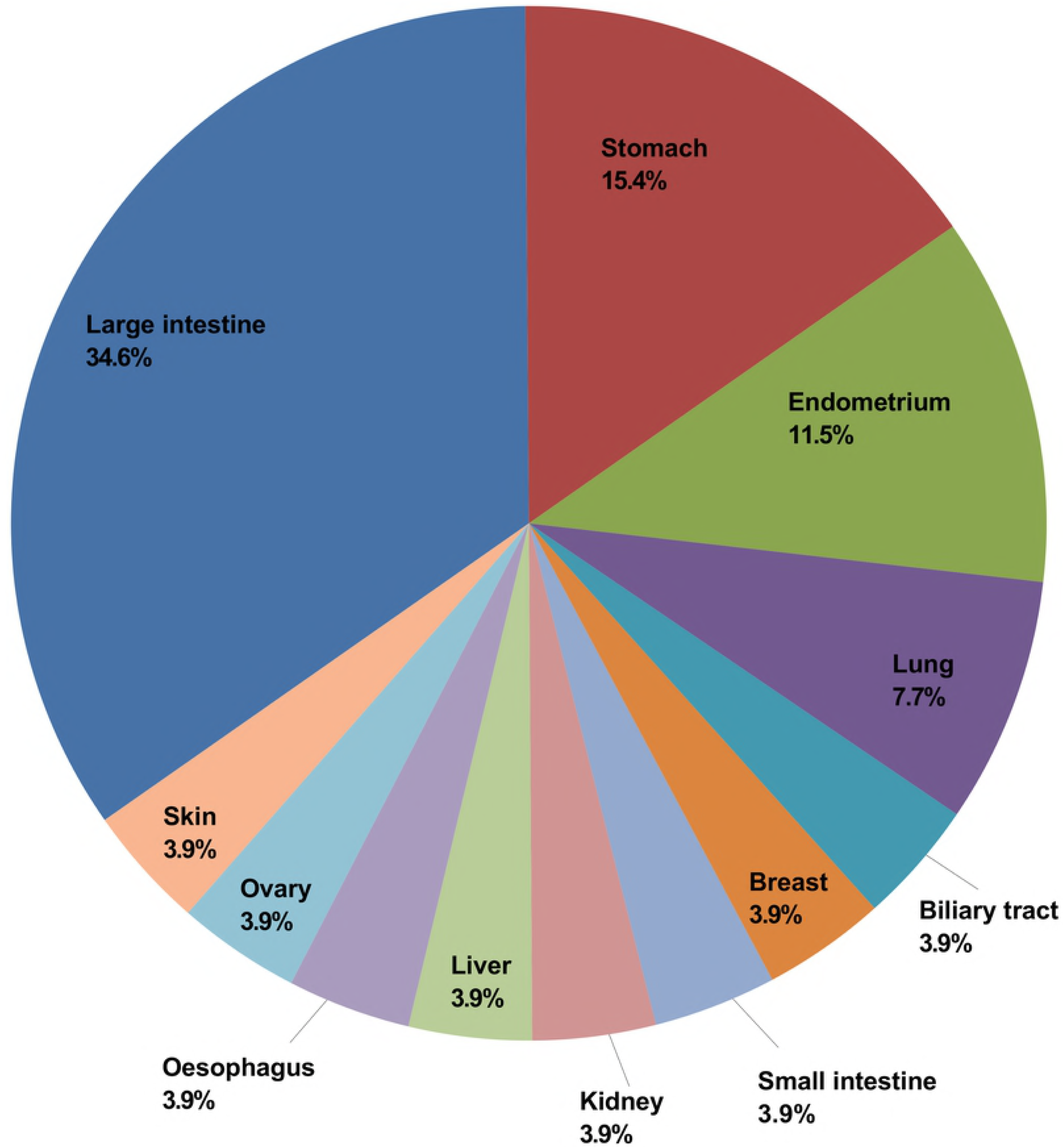
Organs affected by VLP missense mutations in human cancer. Fractions are shown in a single pie chart. Notice the prevalence of intestinal cancers.

## Discussion

Oxygen sensing in mammal cell occurs at many levels, yielding both acute and chronic adaptation. The pVHL/HIF-1α axis is deputed to this role through fine regulation of several hypoxia responsive elements. From a biological network point of view, pVHL is a conserved core element, or a master protein, driving important cell functions. Robustness is a property of biological systems. A relevant aspect of robustness is a clear distinction between proteins acting as signals (modulators) and members of the central control (conserved core) [11]. To guarantee system stability, defined as resistance to external aggressions such as mutations and metabolic imbalance, the central core elements frequently evolve specialized isoforms with partially overlapping functions. This concept is well known for the combined functions of p53 protein family [17]. A similar assumption of functional family can be hypothesized for pVHL. Currently, three isoforms and a paralog protein are described in human for this important tumor suppressor [7,15,20]. The two major isoforms, i.e. pVHL19 and pVHL30, show different cellular localization and the exact nature of their functional segregation remains far from understood. Fewer molecular details are available for VLP. It is thought not to possess ubiquitin ligase activity, missing the pVHL VCB-forming α-domain [53]. This clearly excludes VLP from being a ubiquitin E3 ligase. Our analysis showed this protein is mainly found in primates, where it apparently evolved structural features posing it at middle between pVHL19 and pVHL30. Human VLP is found expressed in three specific tissues characterized by relevant susceptibility to hypoxic injury. The brain, in particular, is well known for its sensibility to variation of oxygen concentration. VLP was proposed as a dominant-negative-VHL exerting a HIF-1α protective function [19]. It is known that neuron-specific inactivation of HIF-1α correlates with increased brain injury during transient focal cerebral ischemia in mice models [64]. In the same study, was also observed that following an initial hypoxic stimulus, HIF-1α levels remained elevated after eight days from a normal oxygenated blood perfusion. Our MD simulations suggest that VLP forms stronger interactions with hydroxylated than unmodified HIF-1α. We believe this data in particular may suggest a precise role for VLP in brain tissue remodeling after a pathological hypoxia insult. In this context, a putative interaction with EP300 (E1A binding protein p300) is worth mentioning. In severe anoxic conditions, p53 outcompetes HIF-1α for EP300 [65], promoting pVHL-independent HIF-1α degradation and leading to a demodulation of hypoxia response. VLP may play a role regulating the competitive p53/ HIF-1α interaction with EP300. Our analysis also sheds light on HIF-1α-independent functions, in particular the putative interaction with eEF1A (Elongation factor 1- alpha 1). This protein is functionally connected with the testis-specific protein Y-encoded (TSPY1) [66], a male specific protein encoded on the Y chromosome expressed in testicular tissue and correlated with spermatogenesis and gonadoblastoma insurgence. As eEF1A mediates pVHL nuclear exporting [67], our investigations suggest the same mechanism to be present in VLP. Combining these findings with VLP expression in testis, we suggest this protein to have a novel regulative role modulating eEF1A/TSPY1 interaction.

## Conclusion

Our analysis shows that VLP shares structural features with the pVHL oncosuppressor in terms of structure conservation and protein-protein interaction partners. It also suggests that VLP could be a member of a larger pVHL family sharing regulative functions. pVHL is a key component of proteasome mediated degradation system, while VLP clearly miss this function. Nevertheless, it may play a relevant role in HIF-1α stabilization. In particular its propensity to form more stable complex with hydroxylated HIF-1α suggests an active role in the modulation of hypoxia response. VLP appears to be structurally very similar to pVHL. Conservation of functional elements at interfaces B and C, in particular, extends the VLP range of activity to HIF-1α-independent functions. These may occur within a precise hypoxia gradient where a finer modulation of pVHL functions would be desirable as in the case of brain remodeling after ischemia injury. The degree to which these results *in silico* apply to living organism is unclear. Experimental studies are in progress to clarify VLP functions.

## Acknowledgements

The authors are grateful to members of the BioComputingUP group for insightful discussions.

